# Using Dual-Coil TMS-EEG to Probe Bilateral Brain Mechanisms in Healthy Aging and Mild Cognitive Impairment

**DOI:** 10.1101/2024.08.23.609391

**Authors:** Ricardo Morales-Torres, Mariam Hovhannisyan, Olga Lucia Gamboa Arana, Moritz Dannhauer, Margaret L. McAllister, Kenneth Roberts, Yiru Li, Angel V. Peterchev, Marty G. Woldorff, Simon W. Davis

## Abstract

**Background:** A widespread observation in the cognitive neuroscience of aging is that older adults show a more bilateral pattern of task-related brain activation. These observations are based on inherently correlational approaches. The current study represents a targeted assessment of the role of bilaterality using repetitive transcranial magnetic stimulation (rTMS).

**Objective:** We used a novel bilateral TMS-stimulation paradigm, applied to a group of healthy older adults (hOA) and older adults with mild cognitive impairment (MCI), with two aims: First, to elucidate the neurophysiological effects of bilateral neuromodulation, and second to provide insight into the neurophysiological basis of bilateral brain interactions.

**Methods:** Electroencephalography (EEG) was recorded while participants received six forms of transcranial magnetic stimulation (TMS): unilateral and bilateral rTMS trains at an alpha (8 Hz) and beta (18 Hz) frequency, as well as two sham conditions (unilateral, bilateral) mimicking the sounds of TMS.

**Results:** First, time-frequency analyses of oscillatory power induced by TMS revealed that unilateral beta rTMS elicited rhythmic entrainment of cortical oscillations at the same beta-band frequency. Second, both bilateral alpha and bilateral beta stimulation induced a widespread *reduction* of alpha power. Lastly, healthy older adults showed greater TMS-related reductions in alpha power in response to bilateral rTMS compared to the MCI cohort.

**Conclusion:** Overall, these results demonstrate frequency-specific responses to bilateral rTMS in the aging brain, and provide support for inhibitory models of hemispheric interaction across multiple frequency bands.

## Introduction

Aging is associated with a number of generalizable changes in brain physiology and function. One of the most common patterns is the increase in bilateral prefrontal activity in response to cognitive tasks, specifically those involving attention or memory [Daselaar and Cabeza, 2005]. The age-related increase in bilaterality in prefrontal cortices (PFC) has sparked a heated debate in recent years [Davis and Cabeza, 2015]. Increases in PFC bilaterality have been interpreted as both a loss of the ability to inhibit the non-dominant hemisphere, with a negative impact on performance [Duverne et al., 2008; Persson et al., 2006], or as a functional compensation, such that the right hemisphere compensates for functions supported by the contralateral cortex in healthy older adults [Davis et al., 2012; Davis et al., 2018]. There is evidence that greater PFC interactivity supports task performance during difficult tasks [Davis and Cabeza, 2015], supporting the latter argument, but data on this phenomenon is sparse, as our understanding of bilateral cortical interactions is limited largely to neuroimaging-based information, which lacks the temporal resolution and causal intervention to make strong claims about cross-hemispheric communication. Non-invasive neuromodulation, coupled with the high temporal resolution of EEG, offers a unique opportunity to address these issues, and to probe the underlying dynamics of hemispheres that need to operate in sync. The current study aimed to address the potential for modulating cross-hemispheric interactions using a novel dual-coil repetitive TMS paradigm and EEG measures of neuroelectric activity.

One major line of thought is that neural processing responsible for bilateral patterns of brain activity is driven by oscillatory patterns at specific frequency bands. Extensive data supports the idea that synchronized neuronal oscillations play a vital role in coordinating interactions between different brain areas [Avena-Koenigsberger et al., 2018; Bressler and Menon, 2010; Varela et al., 2001]. Oscillatory power in the alpha (7–13 Hz) EEG frequency band appears in particular to support long-distance cortical interactions and would be a likely candidate for large-scale cognitive integration across the hemispheres [Chapeton et al., 2019; Lobier et al., 2018]. Synchronized neuronal oscillations could serve a binding role through coordination and regulation of both local processing and inter-areal interactions. As the amount of resources allocated to memory encoding and attentional needs increases, alpha-band power tends to decrease [Fell and Axmacher, 2011; Gevins et al., 1997; Sederberg et al., 2003]. Alpha-band power also tends to increase with age and reduced attentional resources [Scally et al., 2018], as well as with age-related cognitive disorders like Alzheimer’s disease and prodromal forms known as Mild Cognitive Impairment (MCI) [Babiloni et al., 2009; Montez et al., 2009]. Thus, using neuromodulation to affect the expression of this neuroelectric activation frequency in older adults may aid in developing a biomarker of age-related changes in neuroplasticity. Another potential candidate for large-scale cognitive integration within the prefrontal cortices is activity in the beta frequency band (13-30 Hz) [Spitzer and Haegens, 2017]. Long-range beta synchronization has been linked to long-range communication between regions within the frontoparietal attentional network [Kopell et al., 2000] and with the tracking of changes in attentional demand [Gross et al., 2004]. However, the capacity of such entrainment approaches to target and modulate more global communication (bilateral or otherwise) is largely unexplored.

The current study sought to engage a target mechanism (bilateral alpha and/or bilateral beta entrainment) to exogenously modulate cortical oscillations using repetitive TMS (rTMS), thereby establishing the physiological basis for the potential influence of such a mechanism in age-related bilateral recruitment. We hypothesized that while unilateral rTMS would serve to entrain local frequencies, bilateral rTMS would engage global alpha or beta power, commensurate with the view that these frequencies subserve global integration mechanisms. Time-frequency analyses were used to provide high-temporal-resolution information concerning specific electrophysiological responses to rTMS trains in six specific stimulation conditions, using Laterality (unilateral, bilateral), rTMS Frequency (8 Hz, 18 Hz) and Stimulation (active, sham) as factors. The time-frequency analyses were used to analyze the rTMS-train-triggered oscillatory neural interactions between regions. Understanding these neurophysiological activity patterns will help inform optimal choices of stimulation parameters to selectively enhance synchronous communication between left and right PFC in both healthy aging and MCI.

## Methods

### Participants

Twenty-nine individuals participated in this study. Twenty healthy older adults (hOA) were recruited from the community, and nine older adults with Mild Cognitive Impairment (MCI) were recruited from the Duke Neurology Memory Disorders clinic. Twenty-seven participants identified as White/Caucasian and two as Black/African American. Two participants of the 29 did not complete the study due to discomfort from the TMS, and three participants were excluded from data analysis due to extremely noisy EEG data. Thus, a total of 24 participants (18 hOA, 6 MCI) were included in data analysis. Four of the MCI participants were given a formal diagnosis of MCI from a neurologist. Two participants recruited from the community without clinical diagnoses of MCI were categorized as MCI for the analyses based on very low cognitive test scores (MoCA <23). Thus, the MCI group consisted of 6 individuals (3 females; mean age = 71 years, SD = 6.4) and the hOA group of 18 individuals (7 females; mean age = 67 years, SD = 4.2). Participants gave written informed consent and were paid $20 per hour for participating (up to $300). The average time for participation in the rTMS session was about 3 hours. The study was approved by the Duke University Health System Institutional Review Board.

### Screening

Medical history questionnaires were administered to screen participants for TMS and MRI contraindications prior to scheduling. MRI exclusion criteria included, but were not limited to, metallic or magnetic implants, injuries involving metal, and claustrophobia. TMS exclusion criteria included, but were not limited to, history of seizures or epilepsy, neurological disorders, traumatic brain injury, and major psychiatric disorders. Participants completed the Mini International Neuropsychiatric Interview (MINI) to screen for psychiatric disorders. To be eligible, participants needed to be free of neurological or psychiatric medications during the testing period, with the exception of those maintaining a consistent dosage of antidepressants for a minimum of six weeks. Before the first TMS session, a urine drug screen was conducted to rule out undisclosed medications or substances.

### Neuropsychological Assessment

Participants completed The Montreal Cognitive Assessment (MoCA) and the NIH Toolbox Cognition Battery for Age 12+ (NIH Toolbox) to determine their range of cognitive functioning. The NIH Toolbox is an electronically administered assessment consisting of seven subtests assessing executive function, attention, shifting, working memory, episodic memory, language and processing speed [Weintraub et al., 2014]. Performance on subtests considered together is represented in a composite score. Performance on executive function, attention, shifting, working memory, and processing speed subtests provides a fluid intelligence score, whereas language and episodic memory subtests provide a crystallized intelligence score. The MoCA is a widely used cognitive screening tool designed to detect mild cognitive impairment by evaluating various cognitive domains including memory, attention, language, and executive function [Nasreddine et al., 2005]. The MoCA was administered electronically in this study and provides a total score ranging between 1-30, with lower scores indicating more severe cognitive impairment.

Neuropsychological testing scores are displayed in **Table 1**. Due to time constraints of the study and interruptions caused by the onset of the COVID-19 pandemic, MoCA scores were not collected for 5 participants (4 hOA, 1 MCI) at the time of testing. Two of these participants completed the MoCA assessment once research activities resumed. The time between initial data collection and delayed MoCA assessment was 26 and 29 months, respectively for these two participants. One MCI participant did not complete the Picture Sequence Memory subtest of the NIH Toolbox cognitive battery due to fatigue. One hOA participant did not complete the Pattern Comparison subtest due to an administration tablet error. Fluid and composite scores for these two participants were estimated from the remaining seven subtest scores.

**Table 1.**
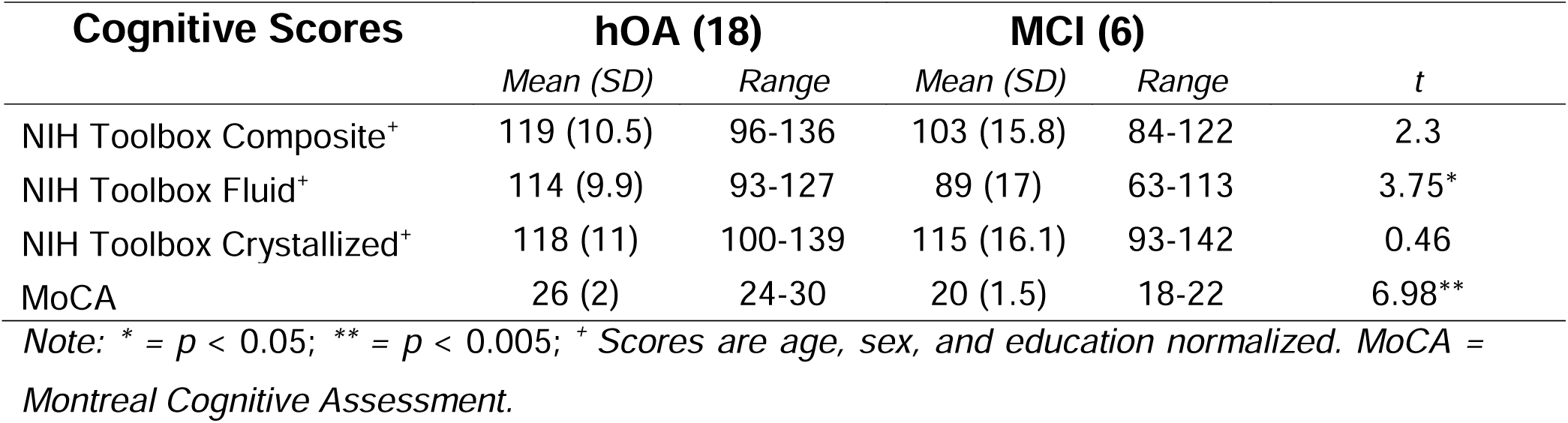
Cognitive Scores.

| Cognitive Scores | hOA (18) |  | MCI (6) |  | *t* |
| --- | --- | --- | --- | --- | --- |
|  | *Mean (SD)* | *Range* | *Mean (SD)* | *Range* |  |
| NIH Toolbox Composite<sup>+</sup> | 119 (10.5) | 96-136 | 103 (15.8) | 84-122 | 2.3 |
| NIH Toolbox Fluid<sup>+</sup> | 114 (9.9) | 93-127 | 89 (17) | 63-113 | 3.75* |
| NIH Toolbox Crystallized<sup>+</sup> | 118 (11) | 100-139 | 115 (16.1) | 93-142 | 0.46 |
| MoCA | 26 (2) | 24-30 | 20 (1.5) | 18-22 | 6.98** |

*Note: \* = $p < 0.05$ ; \*\* = $p < 0.005$ ; <sup>+</sup> Scores are age, sex, and education normalized. MoCA = Montreal Cognitive Assessment.*

### Experimental Design and Equipment

The study consisted of three sessions, each lasting between 2-3 hours. These included (1) an MRI session to collect structural and resting-state imaging and to perform standard neuropsychological testing, (2) a non-task-based simultaneous TMS-EEG session (reported here) during which 6 TMS conditions (described below) were delivered, and (3) a task-based simultaneous TMS-EEG session (not reported here). Sessions always occurred in this order. The current article focuses on the MRI session and the non-task TMS/EEG session. We expand below on the EEG and TMS procedures, then on the MRI procedures.

### EEG Recording

The task protocol was controlled through custom scripts in MATLAB (Mathworks, Natick, MA, USA) using the Psychtoolbox extension (Brainard, 1997) and presented on a 21-inch monitor with 60 Hz refresh rate and 1920 × 1050 resolution. EEG was acquired using a 64-channel actiCHamp EEG system and TMS compatible actiCAP with slim active electrodes (BrainProducts, Gilching, Germany) arranged according to an equidistant, extended-coverage montage [Woldorff et al., 2002]. Electrode impedances were maintained below 5 kΩ, and the EEG data was recorded at a sampling rate of 5000 Hz, with the electrode at *Fz* being used as the online reference. Electric scalp activity was recorded for about 40 minutes in a resting condition with eyes open and focused on a cross-hair fixation point, across several rTMS conditions (**Figure 2B/C**).

### TMS Procedures

Active motor threshold (AMT) was defined as the lowest TMS stimulation intensity required to elicit a motor evoked potential (MEP) when the muscle was actively engaged. Surface electromyogram was recorded from the right first dorsal interosseous muscle by using disposable electrodes (Covidien/Kendall, 133 Foam EMG Electrodes, Mansfield, USA) in a belly tendon montage. After establishing a reliable stimulation site according to the visual-twitch method, AMT was determined by a probabilistic threshold-hunting method using the Motor Threshold Assessment Tool software (https://www.clinicalresearcher.org/software.htm). After finding a defined peak with that site, this area was stimulated 5 times to make sure it was correct before using the threshold-hunting software to determine the AMT. Participants were asked to hold a pressure gauge between their index finger and thumb, and to maintain the needle at about 20% of their total muscle strength. Suprathreshold MEPs were defined as having 50 µV or larger peak-to-peak amplitude within the appropriate temporal range of 25-50 ms after the TMS pulse [Groppa et al., 2012]; MEPs with excessive facilitation or suggestion of a prepotent response or muscle tension were excluded. Stimulation intensity of 80% of the determined AMT was used in the study.

Bilateral rTMS was delivered using a MagPro R30 stimulator and a MagPro X100 stimulator, each connected to a separate Cool-B65 figure-of-eight coil (MagVenture, Farum, Denmark). An Arduino Nano single-board microcontroller was employed to control the triggering of the TMS device to both stimulators. To activate the two MagVenture TMS machines (one R30 and one X100), the Arduino received the rTMS timing parameters via a parallel card. We used *io64* command from Psychtoolbox (Brainard, 1997) to send two specifically chosen byte values (one for each condition: alpha or beta) over the parallel card (PCIe, no hardware interrupts allowed) with a distinct bit set for each value. The sent byte was split using a self-built parallel-to-BNC adapter and connected to two BNC cables one for each distinct bit. These two BNC cables were themselves connected to two separate BNC connectors attached to a self-built housing of the Arduino board. These BNC connectors were connected to two distinct Arduino input pins. Another Arduino pin (connected to the BNC connector on the casing) served as an output trigger and was then split via a coaxial T-junction serving as activation triggers for both TMS devices via a BNC cable for each. The Arduino constantly checked on pin in a continuous loop and sending a trigger signal to both TMS machines accounting for a frequency-dependent delay (alpha: 8 Hz; beta: 18 Hz) with a temporal delay between stimulators of less than 500 µs. Both outgoing trigger cables (connected to one TMS machine each) were digitized with the EEG data via the BrainVision trigger box (**Fig. 1A**). A Brainsight (Rogue Research, Canada) frameless stereotactic neuronavigation system was used to monitor TMS coil position on the participant’s head so that the stimulation location was kept as constant as possible. For this, coil trackers were attached to the coil and the participant’s head, which was registered to the Montreal Neurological Institute standard brain atlas using anatomical landmarks. For the stimulation location, we used standard T1 images to locate the left and right middle frontal gyrus on individual MRIs and marked this location on the EEG cap. A coil holder and a chin rest were also used to maintain stable coil and head positioning. In addition, a spacer was used to minimize contact with the electrodes and associated EEG artifacts (Ruddy et al., 2018), while a thin foam sheet (∼ 1 mm) was added to minimize bone conduction of the TMS sound and scalp sensations caused by mechanical vibration of the coil. Participants wore earplugs for hearing protection and mitigation of auditory activation by the TMS pulse clicks. For the sham conditions, a 3D-printed sham extension was developed to mimic the sound and vibration of the TMS clicks, as well as the appearance of a MagVenture B65 coil (**Fig. 1B**). Two stereotaxic coil holders were used to maintain the coil position over the middle frontal gyrus. Two experimenters were always present in the room, to maintain coil positioning and supervise EEG data quality.

**Figure 1.**
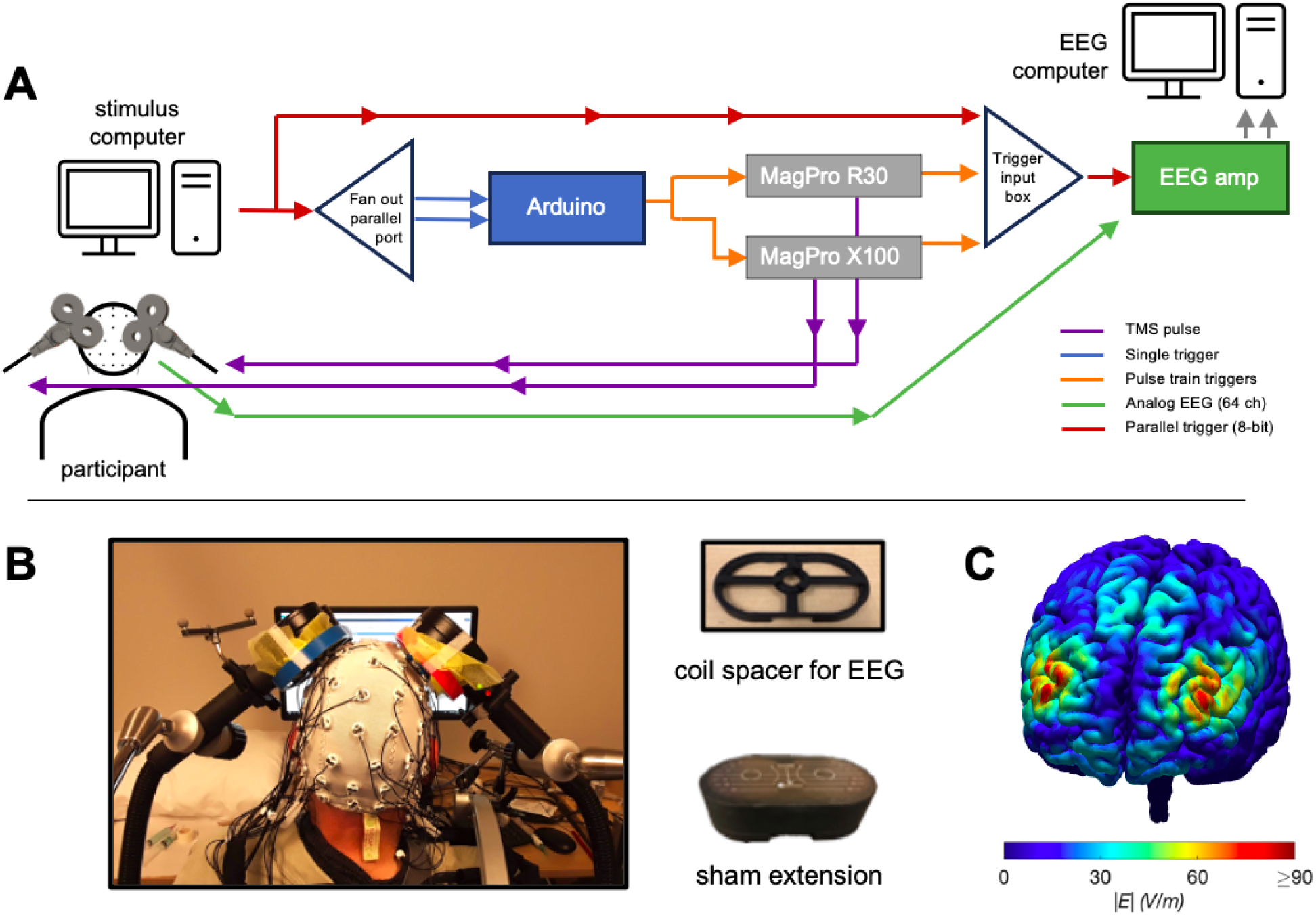
Experimental dual-coil TMS-EEG setup. (A) Wiring diagram. (B) Example setup, with sham extension of the MagVenture B65 coil. (C) Computer model of the magnitude of the electric field |E| induced in the brain by the dual-coil setup at the average pulse intensity in the study simulated in a standard head model (ernie) in SimNIBS 3.2.

Participants were stimulated with 6 different conditions of the rTMS trains: unilateral alpha (8 Hz) (UA), bilateral alpha (BA), unilateral beta (18 Hz) (UB), bilateral beta (BB), unilateral sham (US), bilateral sham (BS). The order of these 6 conditions (**Fig. 2A**) was counterbalanced across participants. Trial-wise timing of rTMS trains (**Fig. 2B/C**) comprised an initial presentation of a fixation cross (at 0 ms), and then 250 ms later the initiation of an rTMS train of 10 pulses delivered at either 8 Hz (alpha band) or 18 Hz (beta band), lasting 1250 or 550 ms, respectively, each at 80% AMT. Trial onsets were spaced 5 seconds apart.

**Figure 2.**
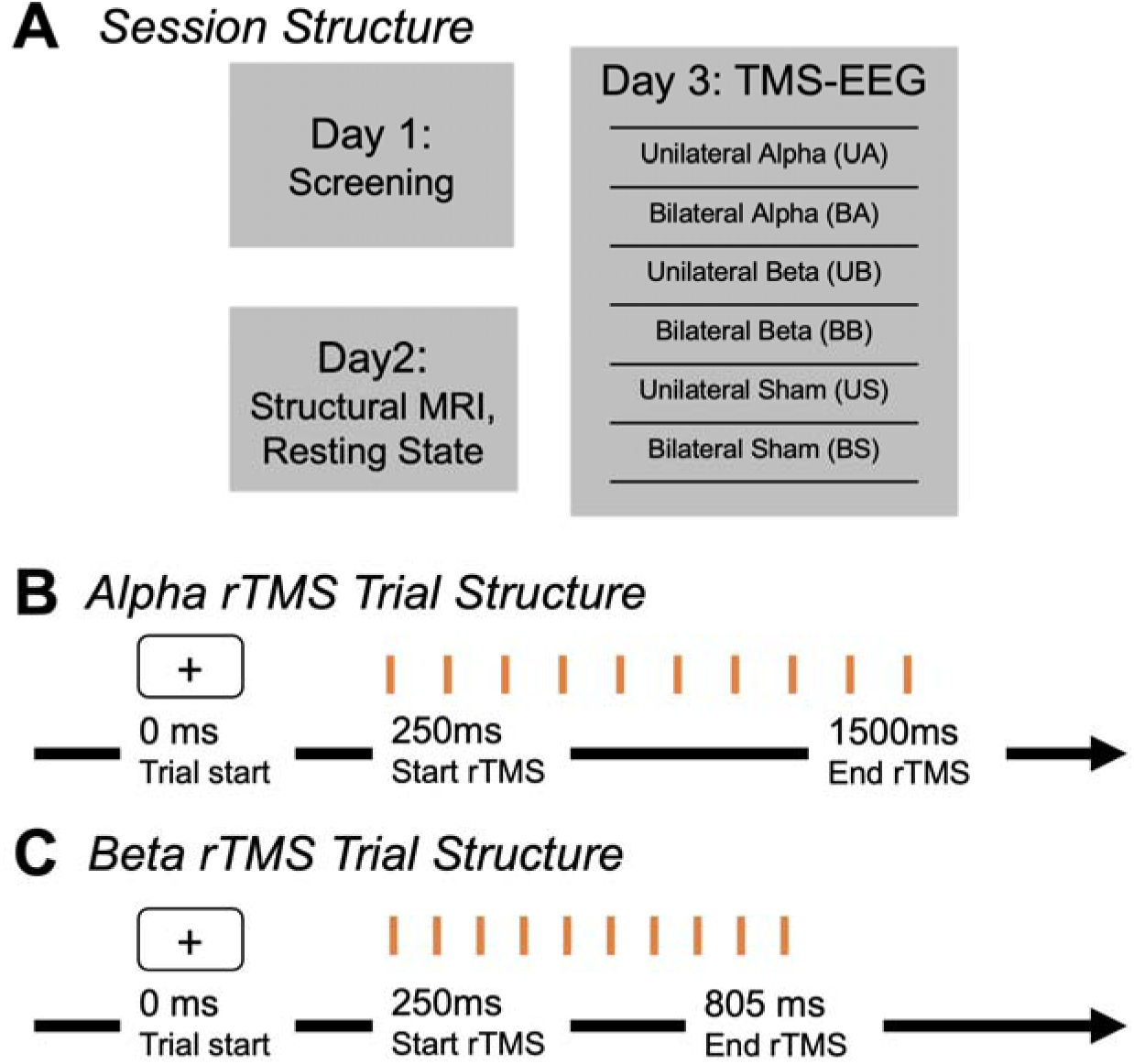
Session and trial structure. **(A)** Session structure comprised Screening, MRI, and rTMS data collection days (plus an additional task-rTMS Day, not reported here). On rTMS days participants were administered rTMS in blocks of 6 different forms of rTMS (counterbalanced orders amongst participants). **(B)** Alpha and **(C)** beta rTMS trial structure; analyses focus on EEG data beginning after the offset of rTMS.

### EEG Analysis

#### EEG Preprocessing

EEG data were preprocessed and analyzed offline using EEGLAB (Delorme and Makeig, 2004) and custom MATLAB scripts (Mathworks, Natick, MA, USA), modeled after Rogasch et al. (2014). Data were down sampled to 1000 Hz and visually inspected to manually remove contamination before further preprocessing. TMS pulses were removed via linear interpolation using data from 10 ms before to 30 ms after each pulse. The EEG signal was bandpass filtered using a zero-phase shift bandpass Butterworth filter (0.1–45 Hz) before applying Independent Component Analysis (ICA) to identify and discard other artifactual components such as eye blinks and muscle activity. ICA-based artifact removal was then performed, leading to an average of 20.5 removed components per participant, with no significant difference in the number of components removed across the 6 TMS conditions (UA, BA, UB, BB, US, BS); researchers were blind to the TMS condition during artifact removal. Data were then down-sampled to 500 Hz, averaged and segmented into epochs. The final analyzed segments from −500 to 3000 ms (relative to the start of a trial, and 250 ms before rTMS) were then re-referenced to the mastoid electrodes (using the algebraic average of the left and right mastoid electrodes).

#### EEG Time Frequency Decomposition

Time-frequency decomposition was performed on the preprocessed single-trial EEG data using a sliding-window short-time Fourier transform with a Hanning window as the window taper. Time-frequency analysis was performed on all artifact–removed trials in each channel, which led to an average of 52.1 trials per condition, per participant. This process generated a time-frequency representation of the data by employing a sliding time window of 500 ms with a 100 ms overlap for each new window, thus corresponding to 400 ms overlap of the sliding windows (i.e., 80% overlap). Activity in frequencies ranging from 1 Hz to 30 Hz were extracted. Power values were then averaged across trials within each condition, transformed into decibels, and referenced to a prestimulus baseline period spanning from −0.9 to −0.4 s before the first pulse (to limit any distortion from spectral spread from the TMS stimulation artifacts).

#### EEG Power Analysis

To compare the six TMS-stimulation conditions, we time-locked the data to the last rTMS pulse in each train and defined two regions of interest (ROIs): the first ROI consisted of the three electrodes beneath the left coil, while the second ROI consisted of the three electrodes beneath the right coil. Subsequently, we computed the average activity for each ROI across four non-overlapping 0.2 s time windows, starting at 0.3 s after the last pulse (again to avoid any potential spectral spread from the TMS stimulation artifact) and extending to 1.1 s. To analyze these data, we conducted a 2 × 4 × 6 mixed-effects model with ROI (Left and Right), Time Window (across the analyzed time range), and rTMS condition (UA, BA, UB, BB, US, BS) as factors, treating each subject as a random effect. These models were implemented using the lme4 R package (Bates et al., 2015), and we utilized the ANOVA function from the *lmerTest* package (Kuznetsova et al., 2017) to identify the best fitting model. Two separate models were performed: the first model examined activity centered on the alpha stimulation frequency (7-9 Hz), while the second model examined the activity centered on the beta stimulation frequency (17-19 Hz).

### MRI Acquisition and Analysis

MRI was performed in a 3-T GE scanner at the at Duke Brain Imaging Analysis Center (BIAC). Structural MRI and diffusion-weighted imaging (DWI) scans were conducted for all participants. The anatomical MRI was acquired using a 3D T1-weighted echo-planar sequence (matrix = 2562, TR = 12 ms, TE = 5 ms, FOV = 24 cm, slices = 68, slice thickness = 1.9 mm, sections = 248). In the fMRI runs, coplanar functional images were acquired using an inverse spiral sequence (64 × 64 matrix, time repetition [TR] = 2000 ms, time echo [TE] = 31 ms, field of view [FOV] = 240 mm, 37 slices, 3.8-mm slice thickness, 254 images). Finally, DWI data were collected using a single-shot echo-planar imaging sequence (TR = 1700 ms, slices = 50, thickness = 2.0 mm, FOV = 256 × 256 mm^2^, matrix size 128 × 128, voxel size = 2 mm^3^, b value = 1000 s/mm^2^, diffusion-sensitizing directions = 36, total images = 960, total scan time = 5 min).

## Results

### Time-Frequency Analyses

Our first analysis looked at the natural peak alpha frequencies across our sample of individuals in order to examine how closely they aligned with our alpha-band stimulation frequency of 8 Hz. To obtain power values that were not influenced by the rTMS, we restricted this analysis to only the sham unilateral and sham bilateral conditions. The peak alpha frequency was determined by averaging the data between −1 s and −0.3 s before the first sham pulse. Subsequently, for each subject, we z-scored the spectrogram in the frequency range of 1 Hz to 30 Hz (**Figure 3**). Across the 24 subjects, the mean natural peak alpha frequency was 8.8 Hz (SD = 1.1 Hz), which thus did align closely with our a priori choice of 8 Hz rTMS. Healthy OA and MCI participants showed no difference in their peak frequency, *t*(7.34) = −0.26, *p* = 0.8).

**Figure 3.**
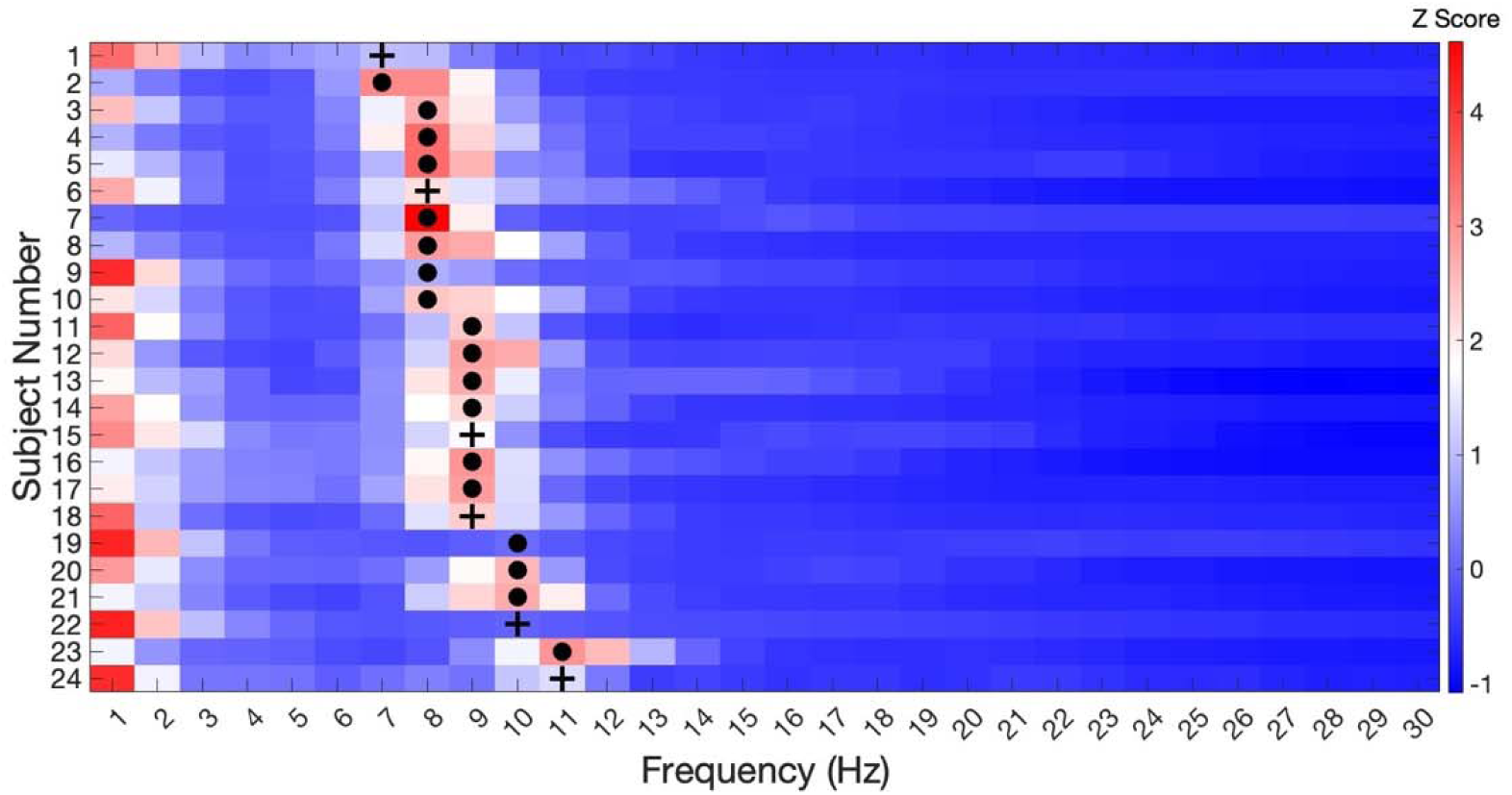
Alpha peaks across subjects. Symbols indicate the peak alpha for each subject. Dots represent healthy older adults while plus signs represent MCI patients.

Moving on to the power analysis, we initially analyzed activity in all participants (hOA + MCI) at the alpha stimulation frequency by averaging power values between 7 and 9 Hz and across the 0.3 to 1.1 s time period (for the data not averaged across the whole time period, see **Fig. 4**). We first tested if the average activity level of each condition was significantly different than 0, with respect to the baseline period. The Bilateral Alpha *(t*(23) = −2.64, *p* = .04) and Bilateral Beta *(t*(23) = −3.22*, p* = .02) conditions were significantly different from 0. The Sham Bilateral *(t*(23) = −1.34, *p* = .38), Unilateral Alpha *(t*(23) = −0.07, *p =* .93), Unilateral Beta *(t*(23) = −052, *p* = .83) and Sham Unilateral *(t*(23) = 0.39, *p* = .83) conditions were not significantly different from 0 (see the upper panel of **Fig. 6**). The False Discovery Rate (FDR) correction was used to reduce the possibility of false positives.

**Figure 4.**
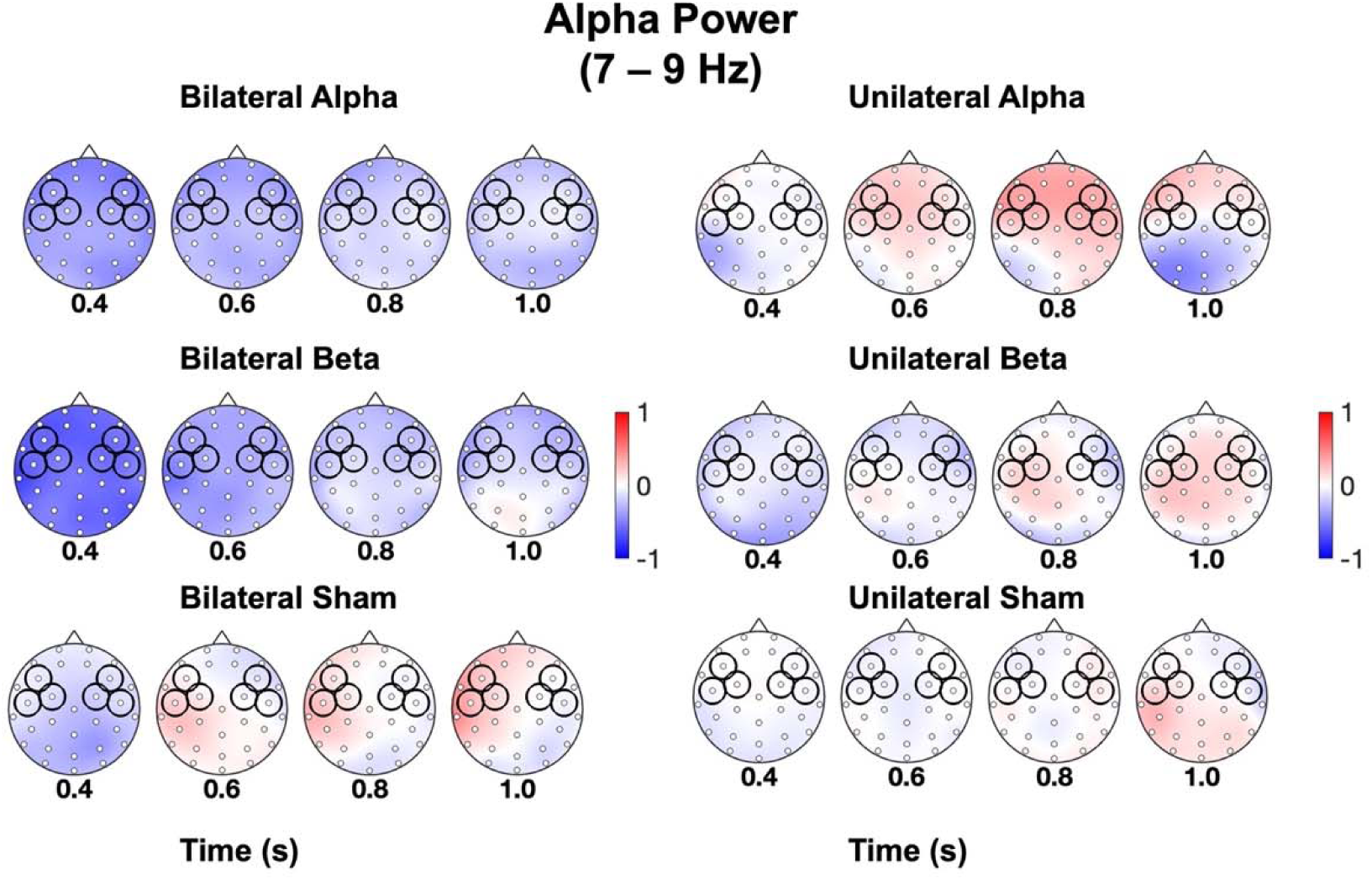
Alpha power associated with unilateral and bilateral rTMS. Time-frequency (TF) plots in the alpha power range (7-9 Hz) shown for 6 rTMS conditions (UA, BA, UB, BB, US, BS), across the topographic map of 64-channel EEG at 4 successive time windows post-stimulation offset (where 0 = offset of rTMS). Power is shown in decibels [−1 1], relative to a prestimulus baseline period spanning from −0.9 to −0.4 s before the first pulse. Left and right electrode ROIs over DLPFC are outlined in black circles.

We then progressed to a more specific comparison to test whether the activity levels in the various conditions varied with the side of recording (Left, Right) and/or timing of stimulation. For this, we employed a model-testing approach, constructing various models from a basic intercept-only model to more complex ones incorporating all variables and interactions. This process determined the best-fitting model for our data. As shown in **Table 2**, the model that best fit included the effects of rTMS Condition and Time, exhibiting a conditional R² of 0.34.

**Table 2.** Model fitting for alpha power.

| Model | Model Fit |  | Model Comparison |  |  |
| --- | --- | --- | --- | --- | --- |
| | AIC | BIC | $\Delta X^2$ | d.f. | p |
| Intercept | 2821.20 | 2836.30 |  |  |  |
| Condition | 2742.10 | 2782.50 | 89.05 | 5.00 | < .001 |
| Condition + Time | 2719.80 | 2775.30 | 28.35 | 3.00 | < .001 |
| Condition + Time + Side | 2721.50 | 2782.10 | 0.25 | 1.00 | 0.61 |
| Condition*Time | 2739.40 | 2875.80 | 12.08 | 15.00 | 0.67 |
| Condition*Time + Condition*Side | 2748.80 | 2910.30 | 0.64 | 5.00 | 0.98 |
| Condition*Side*Time | 2783.10 | 3035.60 | 1.64 | 18.00 | 1.00 |

**Note:** AIC – Akaike Information Criterion; BIC – Bayesian Information Criterion.

Post-hoc pairwise analyses of the best fitting model between conditions revealed that the main effect of condition was driven by the Bilateral Alpha-rTMS condition and the Bilateral Beta-rTMS conditions having lower power in the alpha frequency band, relative to the other conditions. Specifically, the Bilateral Alpha-rTMS condition had lower values compared to the Unilateral Alpha (Estimated Difference = −0.34, SE = 0.1, *p* = 0.002), Sham Unilateral (Estimated Difference = −0.42, SE = 0.1, *p* < 0.001) and Unilateral Beta (Estimated Difference = −0.27, SE = 0.1, *p* = 0.02) conditions. Following the same pattern, the Bilateral Beta-rTMS condition had lower values compared to Unilateral Alpha-rTMS (Estimated Difference = −0.56, SE = 0.1, *p* < .001), Sham Unilateral (Estimated Difference = −0.64, SE = 0.1, *p* < 0.001), Unilateral Beta (Estimated Difference = −0.48, SE = 0.1, *p* < 0.001), and Sham Bilateral Alpha-rTMS (Estimated Difference = −0.37, SE = 0.1, *p* = 0.001) stimulation conditions. All p-values between comparisons are FDR corrected.

The post-hoc analysis also revealed that the main effect of Time was driven by the time window between 0.3 to 0.5 s, which had significantly lower values compared to the 0.7 to 0.9 s (Estimated Difference = −0.2, SE = 0.08, *p* = .03) and 0.9 to 1.1 s (Estimated Difference = −0.32, SE = 0.1, *p* = .001) time windows. All p-values for these comparisons were FDR corrected.

Analogous analyses were performed with the activity in the beta band, for which we averaged activity between 17 to 19 Hz and across the 0.3 to 1.1 s time period (for the data not averaged across the whole time period, see **Fig. 5**). Again, we first tested if the rTMS conditions had a significant effect on the oscillatory power. The Unilateral Beta-rTMS condition was significantly different from 0 (i.e., from baseline) *(t*(23) = 4.01, *p* = .003). The Bilateral Alpha-rTMS *(t*(23) = 0.4, *p* = .9), Bilateral Beta-rTMS *(t*(23) = 0.82, *p* = .83), Sham Bilateral *(t*(23) = 0.12, *p* = .9), Unilateral Alpha-rTMS *(t*(23) = 0.23, *p* = .9) and Sham Unilateral *(t*(23) = 1.3, *p* = .83) conditions did not yield significantly differ from 0 (i.e. the baseline in the beta frequency band (see the lower panel of **Fig. 6**). Again, the FDR correction was used to reduce the possibility of false positives.

**Figure 5.**
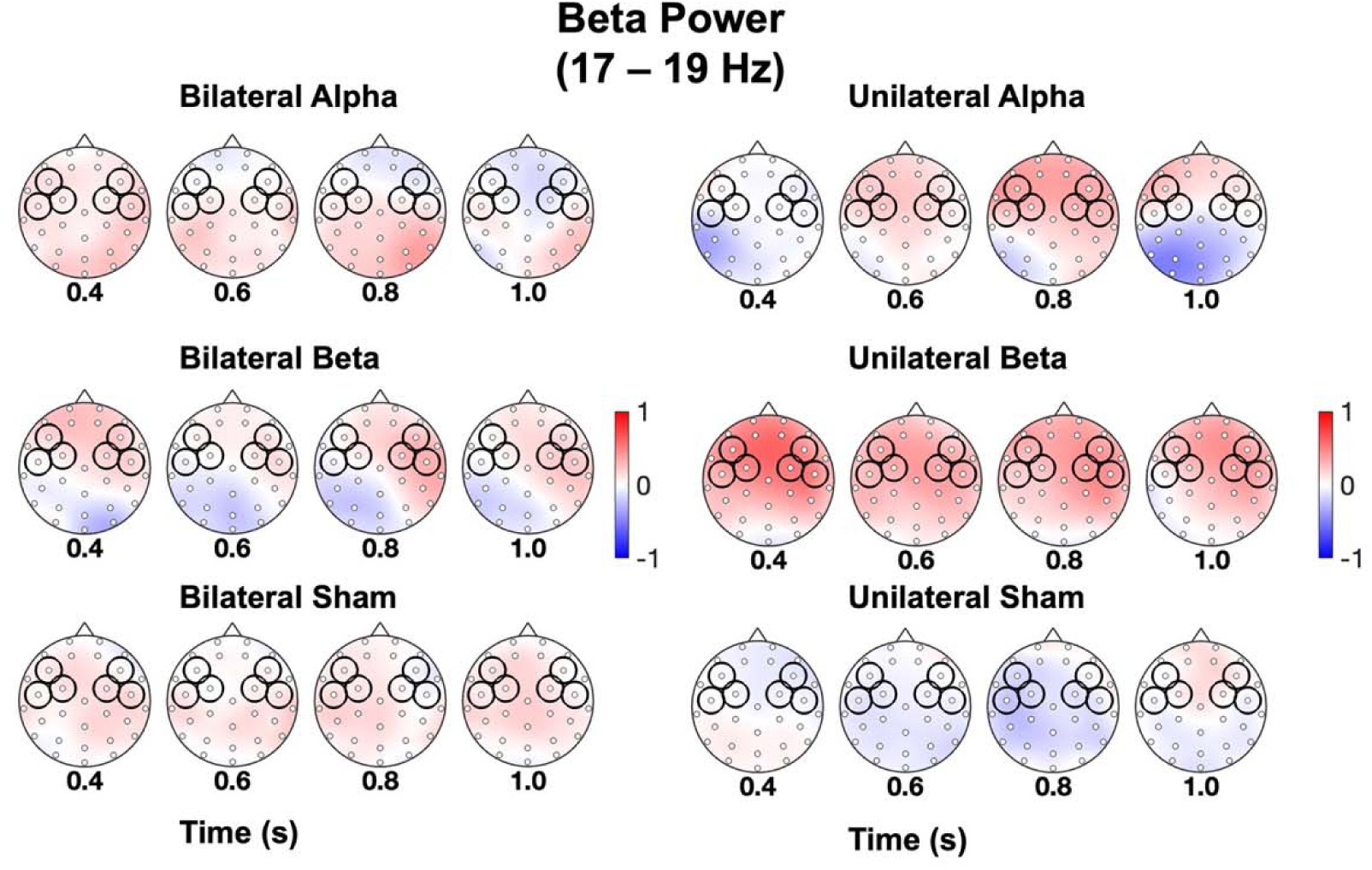
Beta power associated with unilateral and bilateral rTMS. TF results in the beta power range (17-19 Hz) shown for 6 rTMS conditions (UA, BA, UB, BB, US, BS), across the topographic map of 64-channel EEG at 4 successive time windows post-stimulation offset (where 0 = offset of rTMS). Power is shown in decibels [−1 1], relative to a prestimulus baseline period spanning from −0.9 to −0.4 s before the first pulse. Left and right electrode ROIs over DLPFC are outlined in black circles.

**Figure 6.**
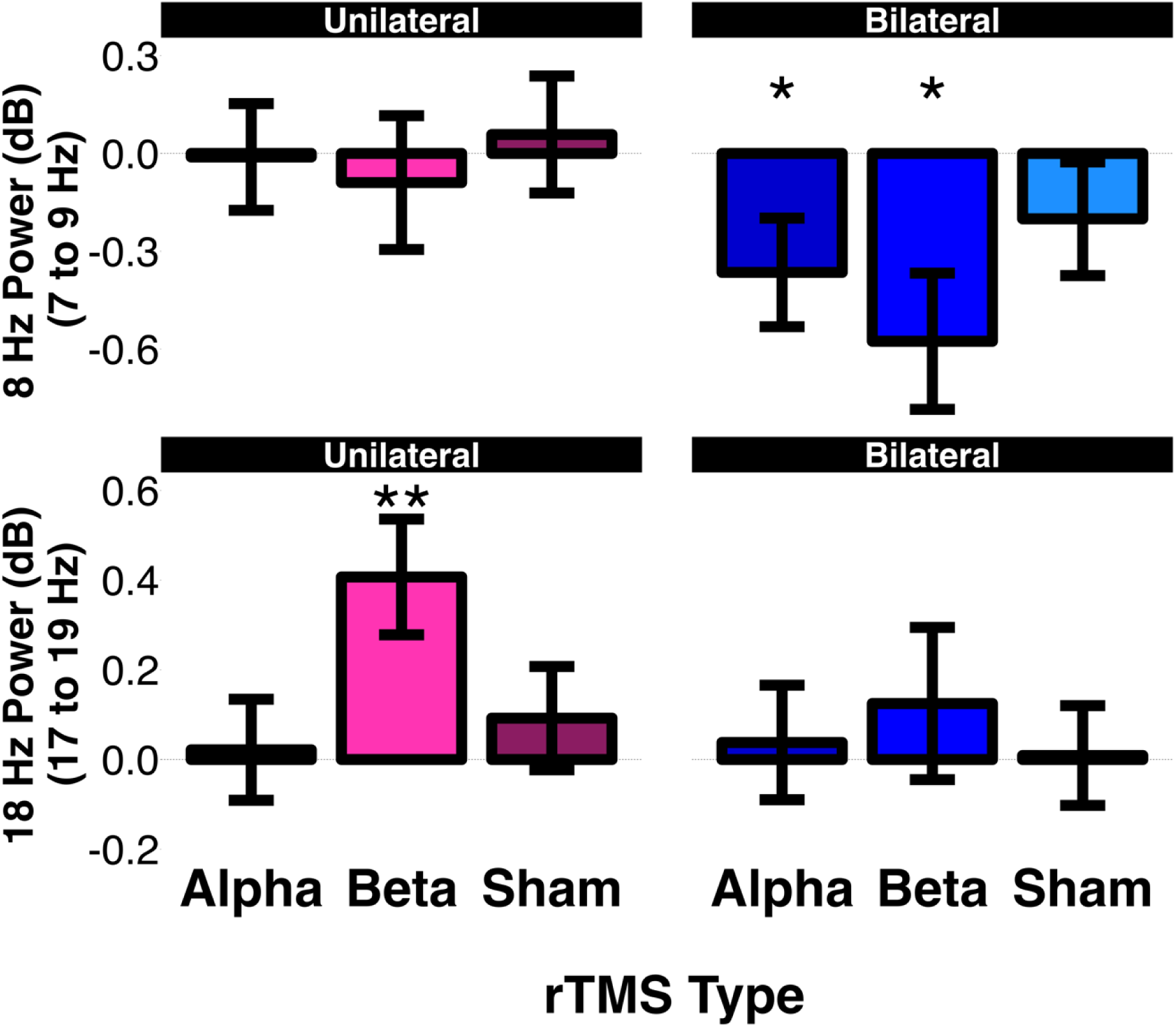
Model estimates of oscillatory power after the last pulse as a function of condition, averaged over the left and right ROIs and over the Time windows (time windows: 0.3 to 1.1 s after last pulse). The upper panel displays the power at 8 Hz (obtained as the average power between 7 to 9 Hz) and the lower panel displays the power at 18 Hz (obtained as the average power between 17 to 19 Hz). Error bars depict the standard error of the mean. * p < .05, *** p < .001.

We then performed the same model-fitting procedure for the beta frequency range that we used for analyzing the activity at the alpha frequency range. As seen in **Table 3**, the best fitting model only included the effect of condition, which shows a conditional R^2^ of 0.17. Post-hoc analysis of this model revealed that the effect of condition was driven by higher beta-band activity values for the Beta-rTMS condition compared with the Bilateral Alpha-rTMS (Estimated Difference = - 0.37, SE = 0.14, *p* = .04), Bilateral Sham (Estimated Difference = −0.39, SE = 0.14, *p* = .04), and Unilateral Alpha-rTMS (Estimated Difference = −0.38, SE = 0.14, *p* = .04) stimulation conditions. All p values between comparisons are FDR corrected. In the supplemental materials, we replicated the analysis using a conventional repeated-measures approach and obtained results consistent with those reported in the above paragraphs.

**Table 3.** Model fitting for beta power.

| Model | Model Fit |  | Model Comparison |  |  |
| --- | --- | --- | --- | --- | --- |
| | AIC | BIC | $\Delta X^2$ | df | p |
| Intercept | 2148.80 | 2164.00 |  |  |  |
| Condition | 2097.60 | 2137.90 | 61.29 | 5.00 | < .001 |
| Condition + Time | 2098.50 | 2154.00 | 5.08 | 3.00 | 0.16 |
| Condition + Time + Side | 2097.80 | 2158.40 | 2.63 | 1.00 | 0.10 |
| Condition*Time | 2120.80 | 2257.10 | 7.05 | 15.00 | 0.95 |
| Condition*Time<br>+ Condition*Side | 2128.10 | 2289.60 | 2.73 | 5.00 | 0.74 |
| Condition*Side*Time | 2162.30 | 2414.80 | 1.71 | 18.00 | 1.00 |

**Note:** AIC – Akaike Information Criterion; BIC – Bayesian Information Criterion.

To address the potential confound of volumetric conduction we re-referenced the data using a Laplacian reference. This type of spatial reference, implemented as the spatial derivative of the local voltage levels, attenuates the effects of volume conduction and provides a more focused view of the local current flows and activity (Cohen, 2015; Kayser & Tenke, 2015). After re-referencing the data, we compared the power across rTMS conditions and frequencies. Consistent with the above-described voltage results, Bilateral Beta rTMS significantly decreased alpha-band activity *(t*(23) = −3.33, *p* = 0.02). Bilateral Alpha rTMS also decreased alpha activity, although this effect was not significant after FDR correction *(t*(23) = −2.32, *p* = 0.09). Unilateral Beta rTMS significantly increased beta power *(t*(23) = 3.75, p = 0.007), suggesting a reliable entrainment effect. Bilateral Beta rTMS also increased beta-band activity, but after FDR correction this effect was no longer significant *(t*(23) = 2.45, p = 0.07). See **Fig. S1**. for the model estimates of each condition.

### Relationship between EEG power modulation and cognition

After establishing the modulation of power induced by the rTMS conditions, we proceeded to analyze whether this effect was associated with individual differences in cognitive status. To do so, we computed the power values averaged across the two ROIs as well as averaged across the five time windows used in the repeated-measures analysis. We then compared these values between the hOA and MCI subjects and subsequently correlated these values with fluid intelligence scores.

We first examined whether there were any differences between hOA and MCI subjects in the alpha power decrease induced by the Bilateral Alpha and Bilateral Beta rTMS conditions. We found that hOA had a significantly lower power at 8 Hz compared to MCI subjects in both the Bilateral Alpha *(t*(14.44) = −3.09, *p* = 0.008) and Bilateral Beta rTMS *(t*(9.70) = −2.68, *p* = 0.02) stimulation conditions (**Fig. 7A**), an effect that was driven by the reduction in power in the healthy older adults (whereas MCI appear unchanged by the bilateral rTMS manipulation). These between-population effects were not seen in the sham conditions. This result suggests that global alpha power is modifiable in our hOA sample, but that this plasticity in alpha reactivity is impaired in MCI participants. In contrast, when examining the power in the beta band induced by Unilateral Beta rTMS, we did not find any significant group difference in evoked beta power *(t*(7.07) =-0.13, *p* = 0.9), suggesting that beta entrainment is feasible in both hOA and MCI populations. Alpha power elicited by unilateral alpha rTMS was not tested, since it did not have a significant entrainment effect in the left PFC (**Fig. 4**).

**Figure. 7.**
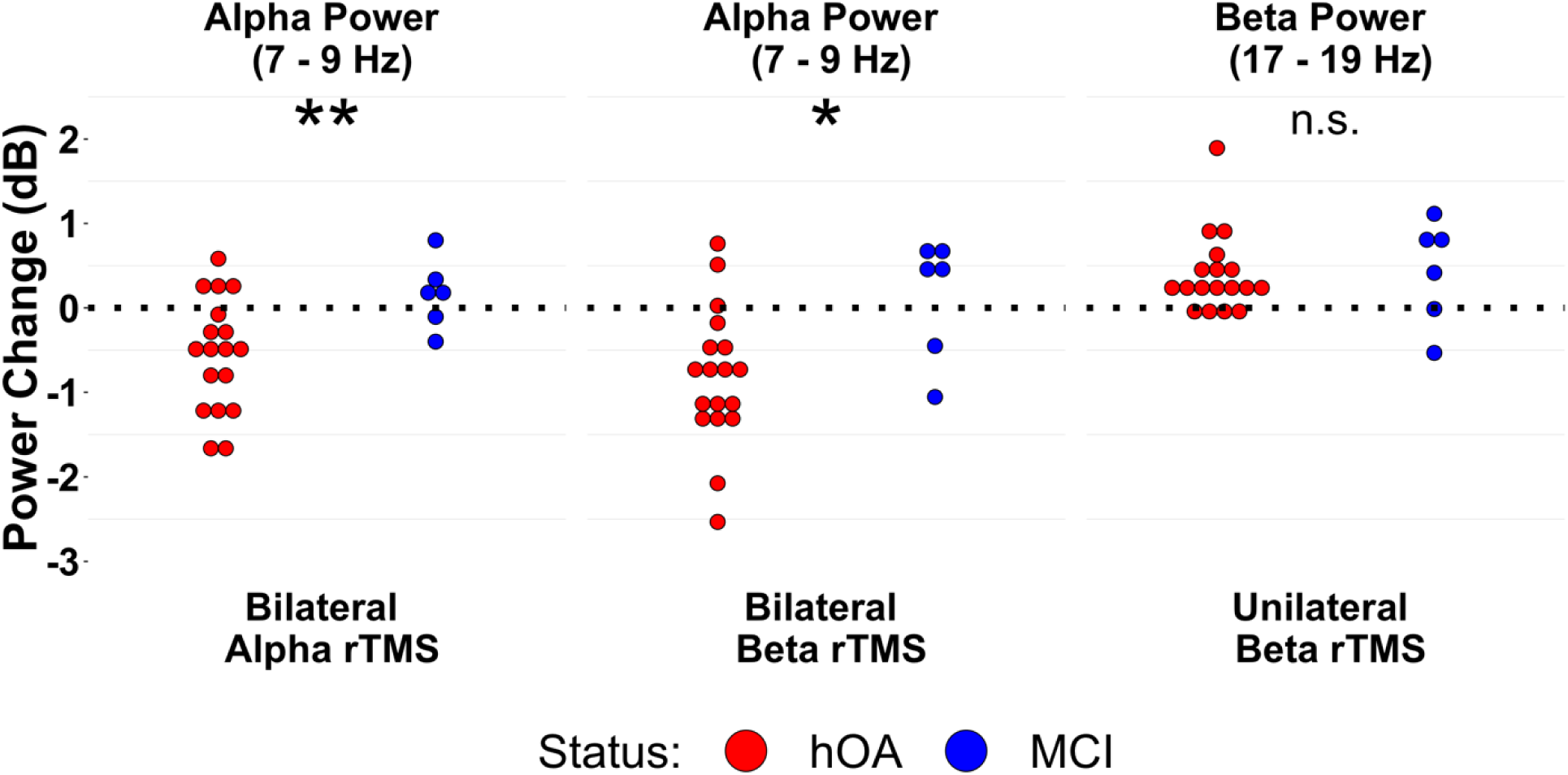
Relationship between Power and MCI Status. The left and center panels show the difference between hOA and MCI in alpha power changes in the Bilateral Alpha and Bilateral Beta stimulation conditions. The right panel shows the difference between hOA and MCI in beta power in the unilateral beta rTMS. * p < .05, ** p < .01.

Turning to the relevance of these patterns for cognition, we then examined the correlation between the rTMS-induced alpha-band power decreases (and beta-band power increases) and fluid intelligence. The activity at 8 Hz was not significantly correlated with fluid intelligence in either the Alpha *(r*(22) = −0.28, *p* = 0.17) or the Beta *(r*(22) = −0.35, *p* = 0.08) Bilateral rTMS conditions. Similarly, Unilateral Beta stimulation was not associated with fluid intelligence, *r* (22) = −0.19, *p* = .36). As such, although there were differential effects on alpha reactivity between the two cognitive groups, it would be difficult to draw any formative conclusions on the specific cognitive import on how beta and alpha power was affected by TMS in our study. However, future TMS-EEG studies employing cognitive tasks known to depend on specific oscillatory frequencies may be more effective at ascertaining such relationships.

## Discussion

A main goal of the present study was to address the neurophysiological dynamics of what may be one of the most fundamental dynamics observed in the aging brain—the increase in bilateral activity in prefrontal cortex in older adults. Time-frequency analyses of post-stimulation effects revealed three main findings, which are generally supportive of an inhibitory — and not linearly-additive — view of hemispheric interactions in a specific frequency band. First, replicating previous unilateral entrainment approaches [Hanslmayr et al., 2014], we found that short trains (10 pulses) of *unilateral* beta (18 Hz) rTMS successfully entrained beta oscillations (i.e., increased beta-band activity). Second, we found that *bilateral* rTMS at both alpha (8 Hz) and beta (18 Hz) stimulation frequencies elicited widespread *decreases* in alpha power. Lastly, we also examined the relevance of these effects for cognition, both by group and continuous measures, and while MCIs showed a weaker bilateral power decrease in the alpha band after bilateral rTMS, we found no qualitative differences in the relationship of these effects and measures of fluid intelligence. We discuss these findings in greater detail below.

Our first main finding was that unilateral bursts of beta frequency rTMS (18 Hz) produced a significant entrainment effect selective to this frequency range (17-19 Hz; see **Fig. 5** and lower left panel of **Fig.6**). Such a result replicates previous demonstrations of beta entrainment; in particular, Hanslmayer et al. 2014 [Hanslmayr et al., 2014] applied 3 separate stimulation frequencies (8, 12, 18 Hz), using a similar 10-pulse rTMS sequence, to entrain cortical oscillations in left prefrontal cortex at those frequencies. In contrast to our unilateral Beta rTMS stimulation effects, Unilateral Alpha rTMS (8 Hz) produced only a weak, non-significant entrainment effect in the PFC for the alpha band (**Fig. 4**), suggesting that this effect may only be successful in cortical regions demonstrating a prominent endogenous frequency in that band, which may not include the PFC. In fact, it is likely that entrainment effects rely on a host of factors [Hanslmayr et al., 2019], including whether the dominant endogenous frequency is present as a result of the brain state of the individual at the time of stimulation. For example, Thut et al. [Thut et al., 2011] used a similar alpha-focused approach to induce alpha oscillations using alpha-band rTMS bursts tuned to the preferred alpha-frequency in parietal cortex; critically, the power of the observed entrainment depended on the phase of the background alpha rhythm prior to the TMS. This result suggests that entrainment depends on synchronizing the TMS train and the endogenous neural oscillator. Such state-dependency is important because it allows distinguishing task-specific mechanisms directly related to cognitive function from effects in oscillatory frequency bands that may be more epiphenomenal, or at least not functionally relevant for successful performance.

Our second main finding was that in contrast to positive entrainment effects specific to unilateral beta stimulation, both bilateral alpha- and bilateral beta-frequency rTMS elicited robust *decreases* in alpha power (upper right panel **Fig 6**). The topology of this effect was widespread (**Fig. 4**). This result has implications for both our understanding of the role of alpha oscillations in coordinating global communication, as well as our understanding of hemispheric interactions. An emerging literature has suggested that synchronized oscillations could serve as a cross-area gating mechanism to quickly enable the selective routing of information between brain regions [Chapeton et al., 2019]. Our unilateral and bilateral findings offer some evidence towards a frequency-specific account on how local exogenous stimulation has more widespread effects on interhemispheric interactions. It has been proposed that a fundamental role of inter-hemispheric interaction is to support contrast-enhancing and integrative functions by co-opting the capacities of the two cerebral hemispheres [Carson, 2020]. However, how the hemispheres interact—and what electrophysiological properties mediate that interaction—is still a matter of great debate [Uddin et al., 2023]. The *interhemispheric inhibition* account is used frequently to describe the action of one hemisphere in suppressing or impeding processing in its counterpart, and is often proposed as a putative mechanism to prevent a bilateral cerebrum giving rise to simultaneous and potentially competing outputs. Such views of the interhemispheric interaction, however, are largely driven by observations in primary motor or sensory systems [Daffertshofer et al., 2005; Ferbert et al., 1992], not higher-level cortical regions such as the PFC. An alternative model, first advanced by Kinsbourne, was that interhemispheric functional connectivity serves to balance the activation patterns between the left and right hemispheres [Kinsbourne, 2003]. In contrast to models that purport that callosal projections are either fundamentally “excitatory” or “inhibitory”, Kinsbourne’s model of *interhemispheric balance* proposes that the balance between the hemispheres occurs primarily through the mechanism of crossed-surround inhibition, which stabilizes cross-hemispheric activation through a positive-feedback opponent-processor interaction. As such, the relevance to the current study is that while unilateral stimulation may entrain lateralized cortex (especially when matched to endogenous frequencies), resulting in a local *increase* in the induced activity at the stimulation frequency, bilateral stimulation with simultaneous trains of rTMS to the two hemispheres—regardless of the entrained frequency— *disrupts* interhemispheric balance, resulting in a global reduction in coordinated alpha oscillations.

Our third main finding was that healthy older adults showed greater TMS-related reductions in alpha power in response to bilateral rTMS, compared to a smaller MCI cohort. In contrast, cognitive status did not have a strong influence on the beta entrainment effect. While MCI participants did not show the widespread depression of alpha power in either bilateral stimulation condition (**Fig. 7A**), they nonetheless showed significant beta entrainment (**Fig. 7B**) in the unilateral (left) beta rTMS condition. The first result is consistent with the observation that MCI/AD status is often associated with reductions in alpha power [Hogan et al., 2003]. This result may offer some promise for interventions focused on age-related memory disorders: Because decreases in resting alpha power have been reported to be reliable biomarkers of AD-status [Montez et al., 2009], establishing a reliable means of modulating alpha oscillations would be an attractive clinical target for AD-related populations. In our sample, bilateral rTMS was unsuccessful in reducing alpha power in MCI participants, but in the future several rTMS parameters (intensity, duration, bilateral timing) could be explored to engender this alpha decrease. On the other hand, and in clear contrast to these induced alpha effects, MCI participants were just as likely as hOAs to demonstrate increases in beta oscillations after unilateral 18 Hz rTMS.

Assuming that beta entrainment reflects a relatively local effect (as suggested by Hanslmayr et al. 2014), and that alpha reductions reflect a more global pattern, such a pair of findings is consistent with the idea that AD-related pathologies are associated with reductions in global plasticity and long-range functional interactions. The disconnection hypothesis of AD purports that disrupted synchrony between distant cortical regions upsets the balance between local specialization and global integration in the brain and consequently impairs cognitive ability [Dai and He, 2014; Wang, 2007]. TMS-induced cortical plasticity in the motor system has been shown to be a good predictor of therapeutic outcome for a PFC-based 10 Hz rTMS intervention in AD populations [Brem et al., 2020]. We note, however, that we did find any significant relationships between the TMS induced alpha reductions and different tests of cognition (fluid and crystalized intelligence). Nonetheless, the current study links such diagnostic predictors of treatment outcomes to a reliable observation from the cognitive neuroscience of aging, namely the age-related increase in bilateral activation patterns. Future work engaging the same mechanisms during an active task may help to establish the functional relevance of bilateral alpha power decreases induced by this form of rTMS.

### Limitations and Methodological Considerations

While the current study advances a new paradigm for investigating a widely observed phenomenon in both younger [Davis and Cabeza, 2015] and older adults [Cabeza, 2002], a number of limitations must be acknowledged. First, the current study used an “in-phase modulation” approach in which both left and right TMS coils stimulated at the same time. Future bilateral TMS applications might incorporate timing delays between the initiation of the left and right hemispheric stimulations trains that incorporate both the phase delays and conduction times between the hemispheres (∼30ms, see [Chapeton et al., 2019]). Second, our choice of stimulation frequencies was the same across our sample (8 Hz and 18 Hz), and not individualized for each participant’s endogenous peak frequencies. There is considerable variation in the peak alpha frequency (see **Fig. 3**), suggesting there likely would be value in utilizing the observed endogenous alpha frequencies. Third, though we observed no linear effects over the timecourse of a block, our trial spacing (5 s) may have nonetheless engendered unexpected cumulative effects [Julkunen et al., 2012]. Fourth, the sham stimulation was applied only at one frequency (8 Hz) owing to time constraints; the development of shams are never perfect but always advancing, and future confirmatory work of these effects should incorporate frequency-matched sham stimulation. Lastly, the sample size of the current study was relatively small.

Taken together, this project tested the feasibility of simultaneous, bilateral rTMS, concurrent with EEG, in older adults (healthy OAs and MCI), to test fundamental questions about how the hemispheres of the aging brain respond to exogenous stimulation. We found that while unilateral rTMS may entrain oscillations for specific frequencies, bilateral rTMS elicits suppression of alpha power for both alpha- and beta-frequency stimulation relative to sham. Moreover, MCI patients showed a significant reduction of this alpha power effect. Lastly, given that this study marks the first application of bilateral, simultaneous rTMS trains, the study also demonstrated the technical feasibility and participant safety associated with this novel form of neuromodulation. More generally, these data support the inhibitory model of prefrontal hemispheric interactions, and thus may help to better understand the age-related increase in bilateral prefrontal activity in aging.

## Acknowledgements

Research reported in this publication was supported by the National Institutes on Aging Awards K01AG053539 and R01AG075417. The content is solely the responsibility of the authors and does not necessarily represent the official views of the National Institutes of Health.

