## Supplemental File for "Using Dual-Coil TMS-EEG to Probe Bilateral Brain Mechanisms in Healthy Aging and Mild Cognitive Impairment"

### Supplemental Materials

**Table S1. rmANOVA on Alpha Power**

| **Predictor** | ***df_Num_*** | ***df_Den_*** | ***Epsilon*** | ***F*** | ***p*** | **η^2^_g_** |
| --- | --- | --- | --- | --- | --- | --- |
| Side | 1.00 | 23.00 |  | 0.35 | .561 | .00 |
| Time | 1.41 | 32.35 | 0.47 | 6.59 | .008 | .02 |
| Condition | 5.19 | 119.32 | 1.04 | 3.40 | .007 | .06 |
| Time x Side | 2.68 | 61.61 | 0.89 | 1.99 | .131 | .00 |
| Time x Condition | 8.15 | 187.46 | 0.54 | 1.08 | .376 | .01 |
| Side x Condition | 4.00 | 91.96 | 0.80 | 0.23 | .922 | .00 |
| Time x Side x Condition | 10.61 | 243.92 | 0.71 | 0.89 | .547 | .00 |

***Note:*** *df_Num_* indicates degrees of freedom numerator. *df_Den_* indicates degrees of freedom denominator. Epsilon indicates Huynh-Feldt multiplier for degrees of freedom, *p*-values and degrees of freedom in the table incorporate this correction. η^2^_g_ indicates generalized eta-squared.

To test whether our results were dependent on our use of mixed-effect models, we repeated the same analysis with a more conventional statistical approach. Two 2x5x6 (rm)ANOVA were run, with ROI (Left/Right), Time (0.3 to 0.5/0,5 to 0.7/0.7 to 0.9/0.9 to 1.1 seconds) and TMS-stimulation condition (Bilateral Alpha/Bilateral Beta/ Sham Bilateral/Unilateral Alpha/Unilateral Beta/ Sham Unilateral). The first model was conducted with alpha power as the dependent variable, while the second model was conducted with beta power as the dependent variable (the values were the same as the ones that were employed during the mixed-effect model).

The first rmANOVA revealed a main effect of condition (F(5.19, 119.32) = 3.4, p = .007) and time (F(1.41, 32.35) = 6.59, p = .008) on alpha power (see **Table S1**). The main effect of condition was driven by the Bilateral Beta condition having significantly lower power values compared to the Unilateral Alpha (t(23) = -2.65, p = .01), Unilateral Beta (t(23) = -2.23, p = .4) and Sham Unilateral (t(23) = -3.08, p = .005). The Bilateral Alpha condition had significantly lower power values compared to the Unilateral Alpha (t(23) = -2.34, p = .3) and Sham Unilateral (t(23) = -3.09, p = .005). The main effect of time was driven by the 0.9 to 1.1 time window having higher power values compared to the 0.3 to 05 (t(23) = 3.05, p = .005), 0.5 to 0.7 (t(23) = 2.9, p =.008) and to the 0.7 to 0.9 (t(23) = 3.15, p = .004) time windows.

The second (rm)ANOVA, conducted with Beta power as the dependent variable (see **Table S2**), only revealed a significant main effect of condition (F(5.05, 116.14) = 2.39, p = .04). This effect was driven by the Unilateral Beta condition having higher power compared to the Bilateral alpha (t(23) = 2.72, p = .01), Sham Bilateral (t(23) = 3.26, p = .003), Unilateral Alpha (t(23) = 2.76, p = .01) and Sham Unilateral (t(23) = 2.31, p = .02).

**Table S2. ANOVA results on Beta Power**

| Predictor | *df_Num_* | *df_Den_* | *Epsilon* | *F* | *p* | η^2^_g_ |
| --- | --- | --- | --- | --- | --- | --- |
| **Side** | 1.00 | 23.00 |  | 3.01 | .096 | .00 |
| **Time** | 1.79 | 41.23 | 0.60 | 2.61 | .091 | .00 |
| **Condition** | 5.05 | 116.13 | 1.01 | 2.39 | .042 | .05 |
| **Time x Side** | 2.26 | 52.08 | 0.75 | 0.30 | .771 | .00 |
| **Time x Condition** | 12.26 | 282.09 | 0.82 | 0.68 | .778 | .01 |
| **Side x Condition** | 4.65 | 106.98 | 0.93 | 0.46 | .794 | .00 |
| **Time x Side x Condition** | 13.51 | 310.70 | 0.90 | 0.68 | .792 | .00 |

*Note:* *df_Num_* indicates degrees of freedom numerator. *df_Den_* indicates degrees of freedom denominator. Epsilon indicates Huynh-Feldt multiplier for degrees of freedom, *p*-values and degrees of freedom in the table incorporate this correction. η^2^_g_ indicates generalized eta-squared.

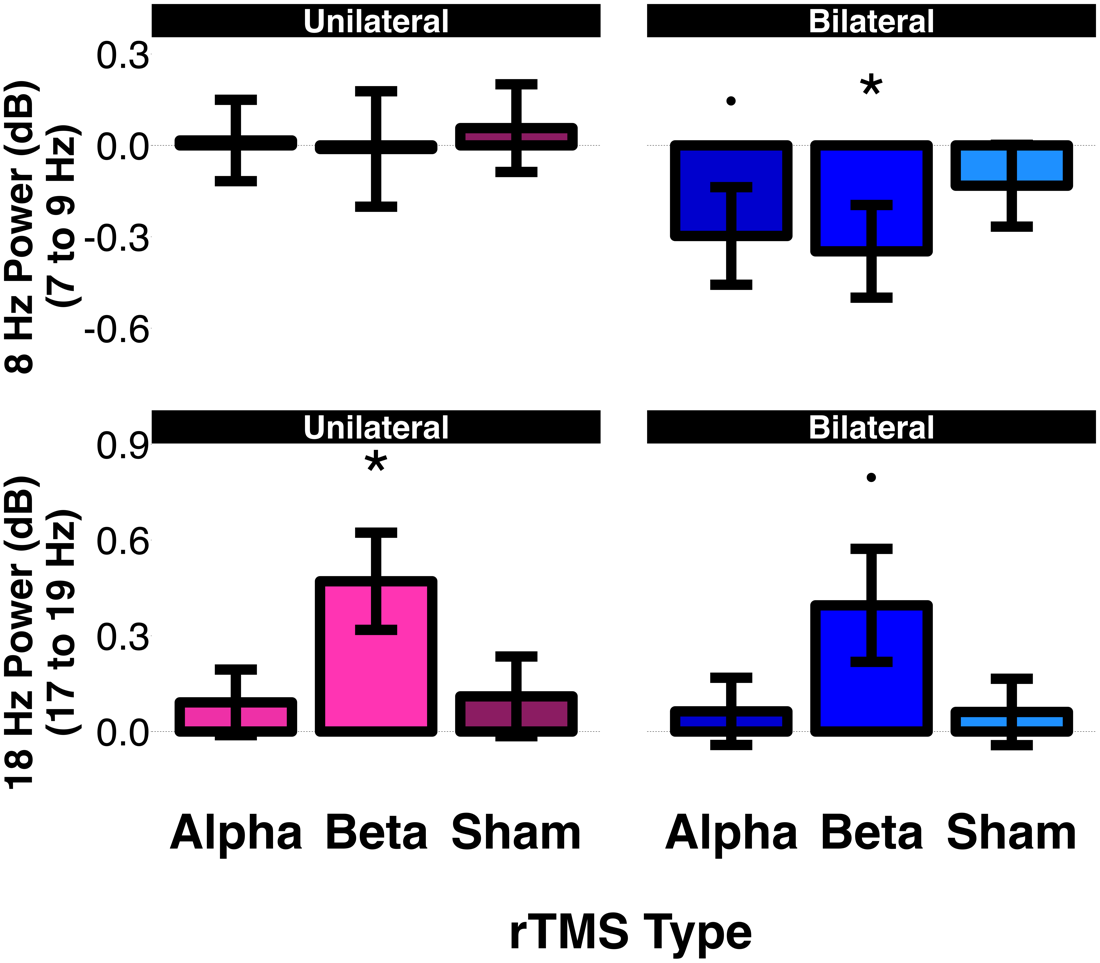

**Figure S.1. Model estimates of the Laplacian re-referenced oscillatory power obtained after re-referencing the data with after the last pulse as a function of condition, averaged over the left and right ROIs and over the Time windows (time windows: 0.3 to 1.1 s after last pulse).** The upper panel displays the power at 8 Hz (obtained as the average power between 7 to 9 Hz) and the lower panel displays the power at 18 Hz (obtained as the average power between 17 to 19 Hz). Error bars depict the standard error of the mean. * p < .05, *** p < .001.
